# A powerful and flexible tool for rapid and accurate differential expression and alternative splicing analysis of RNA-seq data for biologists

**DOI:** 10.1101/656686

**Authors:** Wenbin Guo, Nikoleta Tzioutziou, Gordon Stephen, Iain Milne, Cristiane Calixto, Robbie Waugh, John W. S. Brown, Runxuan Zhang

## Abstract

RNA-seq analysis of gene expression and alternative splicing should be routine and robust but is often a bottleneck for biologists because of reliance on specialized bioinformatics skills. Thus, we have developed “*3D RNA-seq*”, an R shiny App and web based service which provides an easy-to-use, flexible and powerful tool for three-component analysis of RNA-seq data: Differential Expression, Differential Alternative Splicing and Differential Transcript Usage. *3D RNA-seq* integrates state-of-the-art, highly rated differential expression analysis tools and adopts best practice for RNA-seq analysis. It operates through a user-friendly graphical interface, can handle complex experimental designs, allows setting of statistical parameters, tracks results through graphics and tables, and generates figures and a comprehensive report that will guarantee reproducibility. *3D RNA-seq* can be applied to any species and is designed to be run by biologists with no programming skills (or by bioinformaticians) allowing lab scientists to perform rapid and accurate analysis of RNA-seq data.

## Introduction

RNA-seq is generally considered the method of choice to analyse gene expression. The availability of programs such as *Salmon* (Patro et al., 2017) and *Kallisto* (Bray et al., 2016) to quantify transcript and gene level expression accurately and rapidly, allows transcript level analyses to be both feasible and routine. However, the analysis is often a source of frustration for experimental biologists. The key issues are: 1) reliance on the skills of often over-stretched bioinformaticians with the necessary skills to perform the analysis; 2) variation in expertise and knowledge of state-of-the-art methodologies among different bioinformaticians and the continued use of out-of-date and error prone methods/programs (such *Tophat* (D. Kim et al., 2013) and *Cufflinks* (Trapnell et al., 2010)) affecting the quality and reproducibility of analysis; 3) variation in suitability and accuracy of different RNA-seq analysis programs; 4) variation in level of bioinformatics support and access to bioinformaticians’ time (e.g. early career scientists often have more limited access to sufficient bioinformaticians’ time); 5) cost of organisational bioinformatics support; 6) length of time commonly needed (often many months) to generate the data analysis; 7) limitations of many RNA-seq differential expression analysis programs to handle complex experimental designs (e.g. time-course or developmental series data); and 8) despite the ever-increasing appreciation that alternative splicing (AS) is an important level of post-transcriptional regulation, most RNA-seq analyses still focus on the gene level expression thereby ignoring important information that is in the data.

*3D RNA-seq* is an interactive tool for RNA-seq analysis that is implemented using the R Shiny App. It can be run locally or accessed through a web interface (https://3drnaseq.hutton.ac.uk/). It has been developed to carry out 1) differential expression (DE) analysis of genes and transcripts; 2) differential alternative splicing (DAS) and 3) differential transcript usages (DTU) for RNA-seq data, hence *3D RNA-seq*. Isoform switch (IS) analysis for pairwise (Sebestyén et al., 2015) and time-series (Guo et al., 2017) are also incorporated into 3D RNA-seq for advanced AS analysis. The definitions of DE, DAS, DTU and IS are explained in Figure 1. The program integrates the state-of-the-art, highly rated differential expression analysis tool, *Limma* (Law et al., 2014; Ritchie et al., 2015), and adopts current best practice for RNA-seq analysis (Figure 2). *3D RNA-seq* can be used regardless of the type of sample or species under investigation, it can handle complex experimental designs and standardizes the analysis process. An easy-to-use graphical interface takes users through the different steps of the analysis, visualizes the intermediate and final results through graphics and tables, and generates publication quality figures such as heat-maps, volcano plots, histograms, Venn diagrams and expression profiles. Finally, a comprehensive report is automatically generated at the end, which captures all the input info, each parameter used in every step of the analysis, all of the intermediate figures and tables generated as well as the methodologies used at each step. The full analysis report shows the key findings from the analysis and it provides a useful tool to diagnose the technical issues with the RNA-seq experiment and analysis. More importantly, it provides the ultimate guarantee of reproducible analyses. In a typical analysis, RNA-seq data quality control, pre-processing and transcript quantification takes up to two days, and the differential expression analysis and report generation using *3D RNA-seq* takes a few hours (3-Day RNA-seq). There are six main advantages of *3D RNA-seq:* 1) accessibility, ease-of-use, extensive visualization and flexibility allows experimental biologists to control the analysis of their own data; 2) acceleration of RNA-seq analyses – the full analysis can be performed in 0.5 to 2 h depending on the complexity of the experiment; 3) ability to handle complex experimental designs; 4) transcript level analysis for accurate differential expression and differential AS; 5) generation of customised report with details of parameters and figures and tables; and 6) providing consistency and reproducibility in RNA-seq analysis to promote open science.

**Figure 1.**
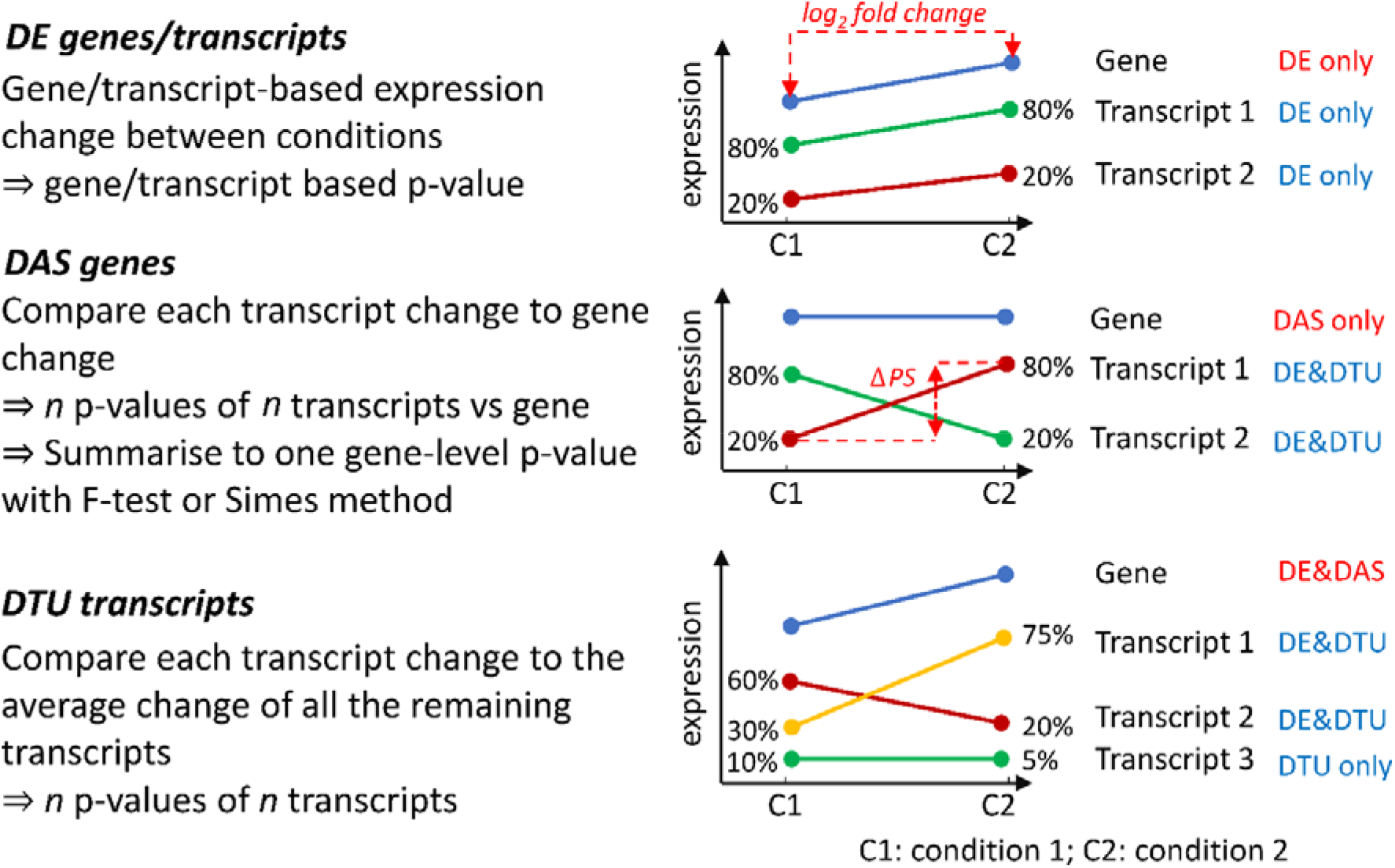
Definitions and criteria for identification of genes and transcripts with significant DE, DAS and DTU. A) DE genes and transcripts are those whose abundance changes between conditions, as measured by changes in log_2_ fold change (L_2_FC). B) DAS genes must have more than one transcript and are determined by comparing the expression changes between individual transcripts to the gene level between conditions. The change in percentage spliced (ΔPS) is calculated as the percentage change in the abundance of a transcript compared to the total expression from gene. For a gene to be DAS, besides a pre-set p-value cut-off, at least one transcript must have a ΔPS ≥0.1. C) DTU transcripts are those transcripts which show different expression behaviour from the other transcripts of the same gene. They are determined by comparing the change in expression of each transcript to the average expression change of all the remaining transcripts of the gene. With these criteria, A) DE only genes are those where the gene and transcript expression levels change significantly but transcripts do not change their relative abundance within a gene. B) DAS only genes are those where the gene expression level does not change significantly but that of at least one transcript changes its relative abundance within a gene. C) DE+DAS genes show both significant gene level expression changes and relative abundance changes of at least one transcript. Isoform switches (ISs) happen when a pair of transcripts reverse their relative abundance across different conditions or time points

**Figure 2.**
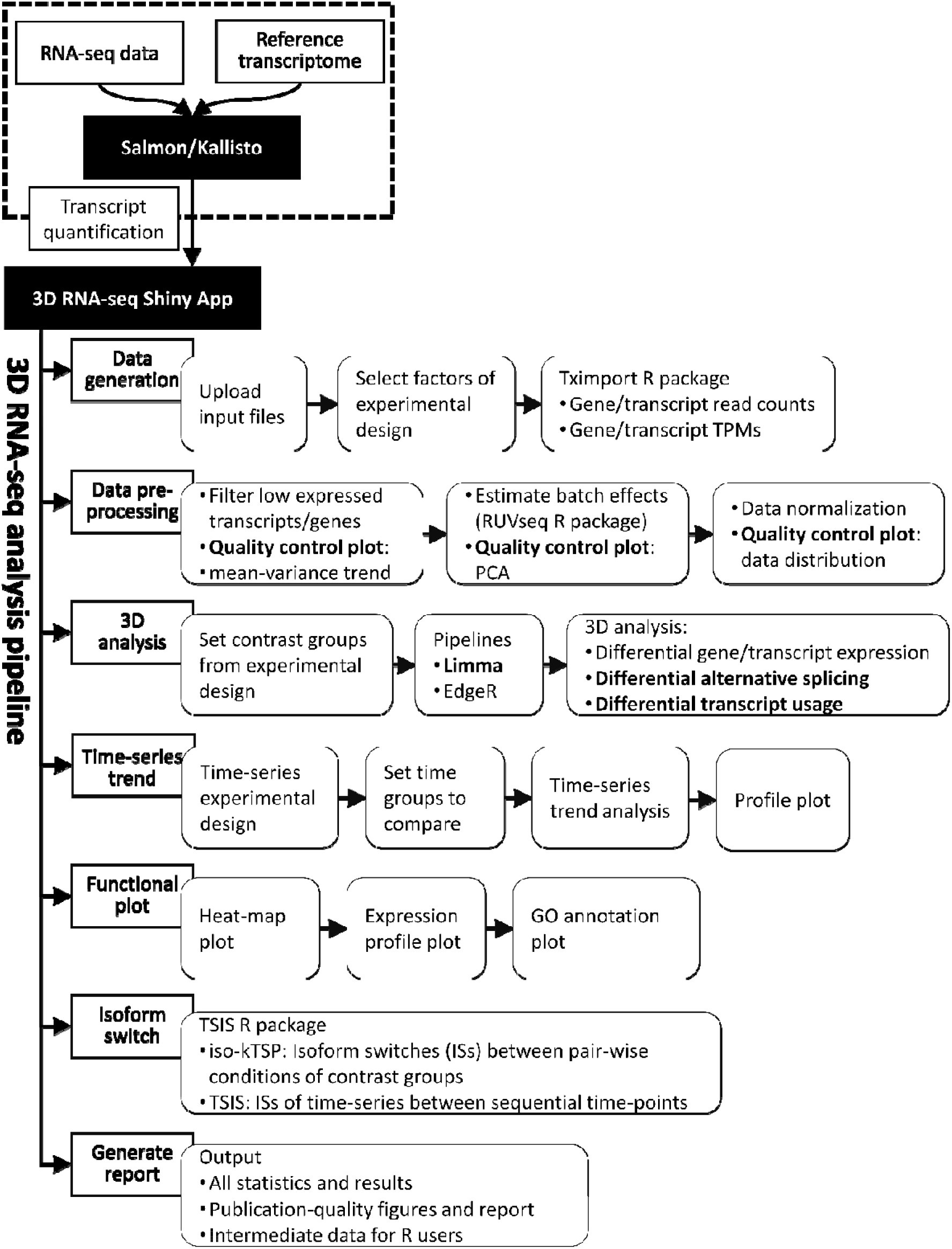
3D RNA-seq analysis pipeline.

## Results

### Application of 3D RNA-seq to RNA-seq analyses of expression/AS in plants

To demonstrate its utility, *3D RNA-seq* was applied to a subset of data from an RNA-seq time-series Arabidopsis plants exposed to cold stress (Additional file 1: Figure S1) (Calixto et al., 2018, 2019). Briefly, 5-week-old Arabidopsis Col-0 plants were grown at 20°C for 24 hours, then the temperature was reduced to 4°C. Samples were harvested every 3 hours for the last day at 20°C, Day 1 at 4°C and Day 4 at 4°C, yielding 26 time-points in total (Additional file 1: Figure S1A). The data from six of these time-points was extracted to illustrate the utility of *3D RNA-seq*. The six time-points are 3 and 6 h into the dark period from the 20°C Day, Day 1 and Day 4 at 4°C (time points T2 and T3, T10 and T11, and T19 and T20, respectively, referred to here as time-points T2 and T3 from Day 0, Day 1 and Day 4 (Additional file 1: Figure S1A). The T2 and T3 time-points represent the equivalent time-point in the three days and thereby control for time-of-day expression variation. Transcript quantification was generated using *Salmon* (Patro et al., 2017) and AtRTD2-QUASI (Zhang et al., 2017).

In the data pre-processing procedure, we removed the low expressed genes/transcripts using the mean-variance trend plots (Additional file 1: Figure S2) (Law et al., 2014) and corrected batch effects using the RUVSeq method (Risso et al., 2014) in the App (Additional file 1: Figure S3). Expression data was normalised across samples to reduce sequencing biases. We also employed stringent filters to control the FDR of multiple testing.

Using *3D RNA-seq*, contrast groups were set up to compare the equivalent time-points before and after cold stress to control for time-of-day variation in expression due to photoperiodic and circadian changes (Calixto et al., 2018). For example, T10 (T2 in day 1 at 4°C) was compared to T2 in day 0 (20°C), T19 (T2 in day 4 at 4°C) was compared to T2 in day 0 (20°C) and so on (Additional file 1: Figure S1B). Other contrast groups can also be set up (Additional file 1: Figure S4). For example, the mean between T10 and T11(T2 and T3 in day 1 at 4°C) can be compared to the mean of T2 and T3 in day 0 (20°C) etc. or the difference between T10 and T11(T2 and T3 in day 1 at 4°C) can be compared to the difference between T2 and T3 in day 0 (20°C) etc. (Additional file 1: Figure S1C). We identified 5,023 DE genes and 2,346 DAS genes. These included 1,875 genes which were regulated only by AS (no significant differential expression at the gene level) and 4,185 DTU transcripts (Figure 3A and B). In addition, 471 of the DAS genes also had significant abundance (DE) changes across the contrast groups (DE+DAS genes). The abundance of significant DTU transcripts can either change significantly (DE+DTU) or non-significantly (DTU-only) (Figure 1). Output histograms illustrate the variation in up- and down-regulation of DE genes and the number of significant isoform switches between the contrast groups (Figure 3C and D). Volcano plots of fold changes in abundance (log_2_FC) against significance (FDR) highlight those DE genes and transcripts with the largest and most significant changes in the data (Figure 3E). Heat maps of clustered DE genes and DTU transcripts are shown for the contrast groups of the six time-points in Figure 4A and B. Functional annotation analysis was performed using RDAVIDWebService R package with significance threshold FDR < 0.05 (Fresno & Fernández, 2013). The DE genes showed GO enrichment in cold-induced physiological and molecular events such as response to various stresses, transcription and altered ribosome production (Figure 4C). Similarly, DAS genes were significantly associated with the spliceosome, RNA splicing and nucleotide/RNA/mRNA binding terms reflecting cold-induced AS of splicing factors (Figure 4D). The Kyoto Encyclopedia of Genes and Genomes (KEGG) pathway enrichment analysis showed DE genes were significantly enriched with Starch and sucrose metabolism, Biosynthesis of secondary metabolites, Plant hormone signal transduction, Ribosome biogenesis in eukaryotes, Photosynthesis – antenna proteins, Plant-pathogen interaction and Glutathione metabolism pathways, while the DAS genes were related to Spliceosome and mRNA surveillance pathways (Additional file 1: Table S1). These results reflect the analysis of the complete time-series (Calixto et al., 2018). Experimental validation of expression, AS and isoform switches by high resolution RT-qPCR and high-resolution RT-PCR have been described previously (Calixto et al., 2018, 2019).

**Figure 3.**
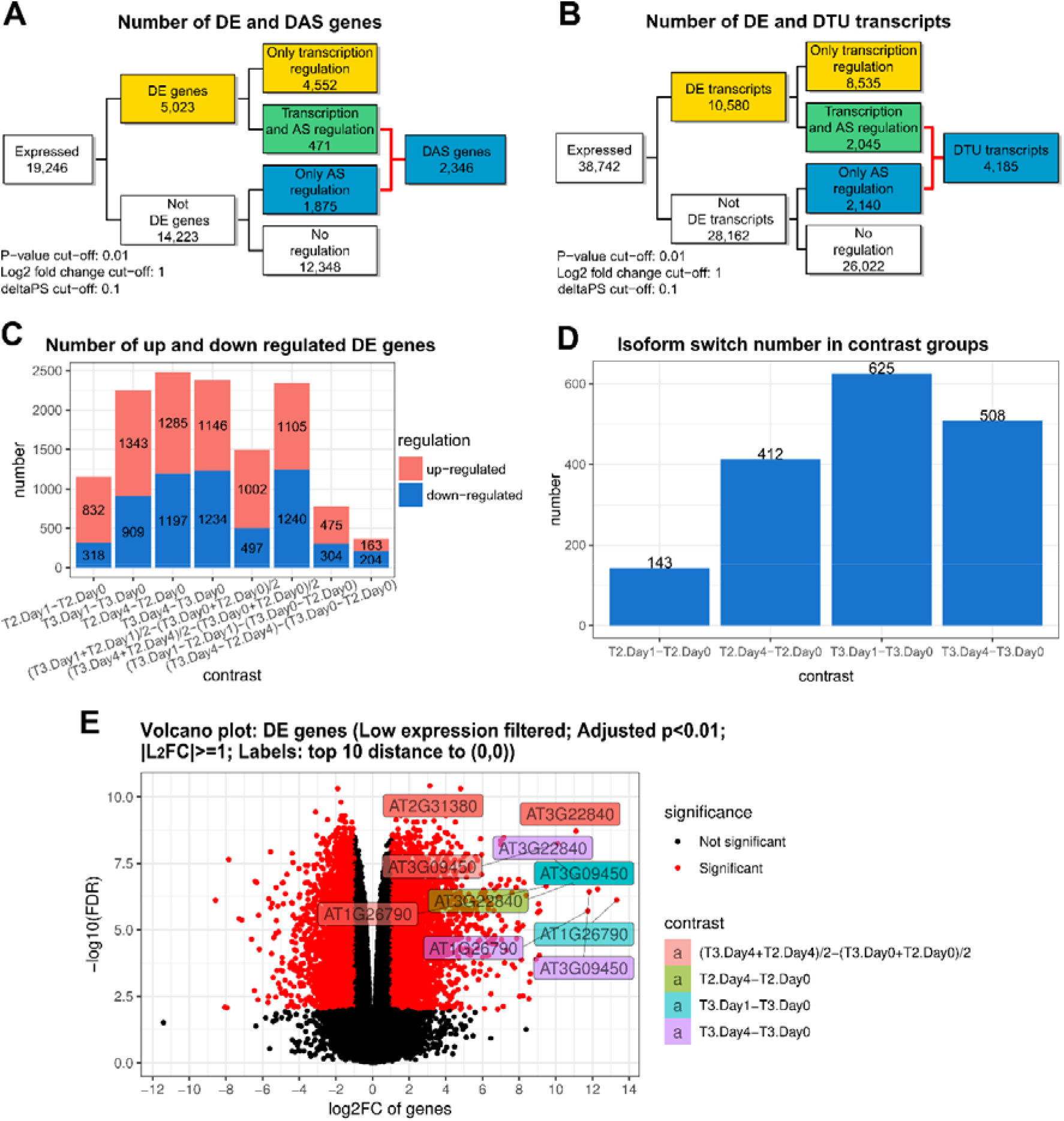
Illustrations of visualization outputs from 3D RNA-seq. A) Summary figure of expressed genes and significant DE, DE+DAS and DAS genes from analysis of the Arabidopsis data; B) Summary figure of expressed transcripts and DE, DE+DTU and DTU transcripts; C) Number of significantly up- and down-regulated DE genes in different contrast groups, and D) Number of significant isoform switches in contrast groups. E) Volcano plot of significant DE genes. The top 10 genes with the smallest *p* values and biggest fold changes are highlighted and different colours refer to different contrast groups.

**Figure 4:**
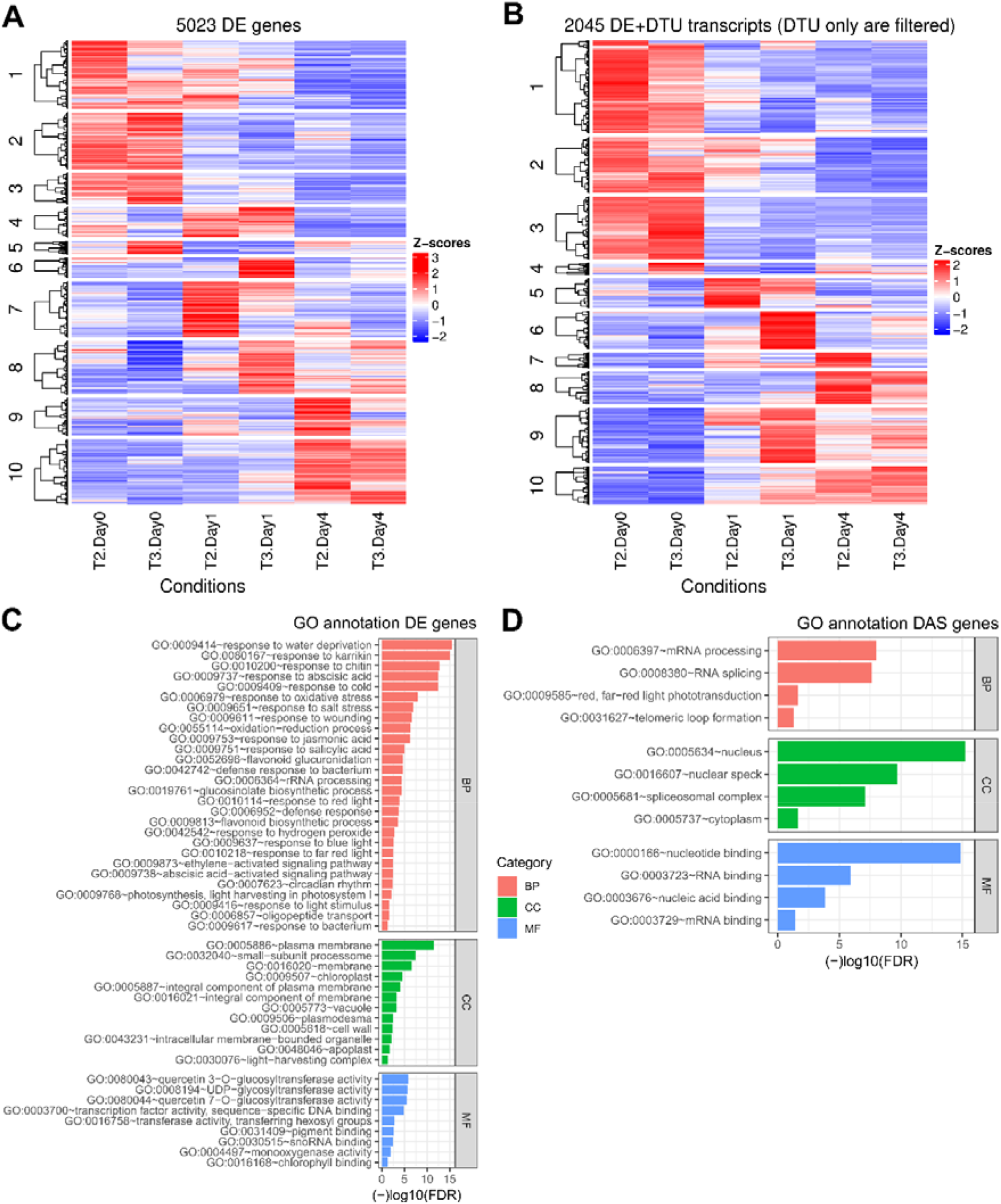
Visualization of clustered genes and transcripts and enriched GO terms. Heatmaps show the grouped expression profiles for A) DE genes and B) DTU transcripts across the samples. The top enriched GO terms for C) DE and D) DAS genes are visualized with their associated FDRs.

The *TSIS* analysis identified 1,688 significant switches between 1,144 pairs of alternatively spliced transcript isoforms in DAS genes with stringent cut-offs: probability > 0.5, sum of average abundance differences > 1 TPM and p-value < 0.05 (Figure 3D) (Guo et al., 2017). These ISs were related to various AS events and approximately half of the isoforms were protein-coding transcripts with different functions according to the transcript annotations in AtRTD2 (Zhang et al., 2017). The switch frequency along the time-series showed that ISs occurred in the contrast groups comparing 4°C and 20°C time-points (Figure 3D). Thus, a large number of genes and transcripts in Arabidopsis are sensitive to temperature reduction and likely contribute to acclimation and survival through AS regulation.

### Application to RNA-seq analyses in animals

To illustrate the utility of the *3D RNA-seq* App in analysing multi-factor RNA-seq data from animals, we re-analysed RNA-seq data which studied the effects of dexamethasone treatment on cortical and hypothalamic neural progenitor/stem cells (NPSCs) from male and female mice (Frahm et al., 2018). The RNA-seq data consisted of male and female cortex and hypothalamus cell cultures treated or untreated with dexamethasone, each with three biological replicates (24 samples in total). The study identified differentially expressed genes common and unique to brain region, gender and dexamethasone treatment (Frahm et al., 2018). The same dataset has also been used to demonstrate the improved resolution of differentially expressed genes using *Sleuth* for transcript-level differential analysis and aggregation of transcript level *p*-values to give gene-level *p*-values (Yi et al., 2018). Using this data, we performed two differential expression comparisons between: 1) the *Sleuth*/aggregated *p*-values method (Pimentel et al., 2017; Yi et al., 2018) and *3D RNA-seq* and 2) the results of the differential analysis in Frahm *et al*., (2018) and those generated by *3D RNA-seq* expression results.

To compare the results from *3D RNA-seq* and *Sleuth* directly, the *Kallisto* transcript quantifications were downloaded (see Availability of data and materials) and pre-processed in *3D RNA-seq*. This identified 43,836 expressed transcripts which had at least 3 samples with ≥ 1 CPM and 12,155 expressed genes which had at least one expressed transcript which were then used in the *Sleuth* and *3D RNA-seq* analyses. In addition, the *Sleuth* analysis in Yi *et al*., (Yi et al., 2018) only examined the effects of dexamethasone and did not distinguish brain region or gender effects and therefore the same contrast groups were set up in *3D RNA-seq* to match the analysis in Yi *et al*., (Yi et al., 2018). We then ran the *Sleuth*/aggregated *p*-values pipeline (see Availability of data and materials) and *3D RNA-seq* on the 43,836 expressed transcripts. Using *Sleuth*, we identified 3,237 DE genes (Figure 5). GO enrichment analysis was performed using Fisher’s exact test in topGO R package (Alexa & Rahnenfuhrer, 2019) in conjunction with genome annotation for mouse org.Mm.eg.db (Carlson, 2019). KEGG enrichment was queried against the KEGG database *(KEGG PATHWAY Database*, n.d.) by using Fisher’s exact test. Significantly enriched GO terms in categories of biological process (BP), cellular component (CC) and molecular function (MF) and KEGG pathways were determined with FDR < 0.05. Significant enrichment terms and pathways relevant to response to stress, immune system, inflammation, hormone response, splicing/spliceosome were extracted to illustrate effects of dexamethasone treatment on gene function (Figure 6; Additional file 1: Table S2). The *3D RNA-seq* analysis used the same contrast groups and a BH adjusted *p*-value cut-off of < 0.05 to identify significant changes (Y Benjamini & Hochberg, 1995). Note that to maintain similar parameters for the direct comparison to *Sleuth*, the log_2_ fold change and Δ percent spliced cut-offs were not applied in the *3D RNA-seq* analysis. The results of the *3D RNA-seq* pipeline showed a high degree of similarity with the *Sleuth* pipeline in terms of identified DE genes but in addition resolved differentially alternatively spliced genes from DE genes and identified 1,649 genes with significant expression/AS changes. *3D RNA-seq* identified a total of 4,284 genes with differential expression and/or differential alternative splicing of which 3,700 and 896 were DE and DAS genes, respectively (with an overlap of 312 gene – DE+DAS genes) (Figure 5). The *Sleuth* pipeline (Yi et al., 2018), which does not identify DAS genes, recovered 3,237 DE genes. Of the total (DE and DAS genes) and DE genes identified by *3D RNA-seq*, 2,635 and 2,346, respectively, were common to both analyses such that 81.4% and 72.5% of the total and DE genes identified by *Sleuth* were identified by *3D RNA-seq* (Figure 5). The *3D RNA-seq* pipeline also identified 5,573 DE transcripts and 1,480 DTU transcripts with adjusted *p*-value < 0.05. Interestingly, 531 of the DAS genes were among the DE genes defined by *Sleuth* – of these, 242 were DE+DAS. Of the 602 DE genes unique to the *Sleuth* analysis, 372 had significant DE/DTU transcripts in *3D RNA-seq* and did not carry over to significant DE or DAS genes (Figure 5).

**Figure 5.**
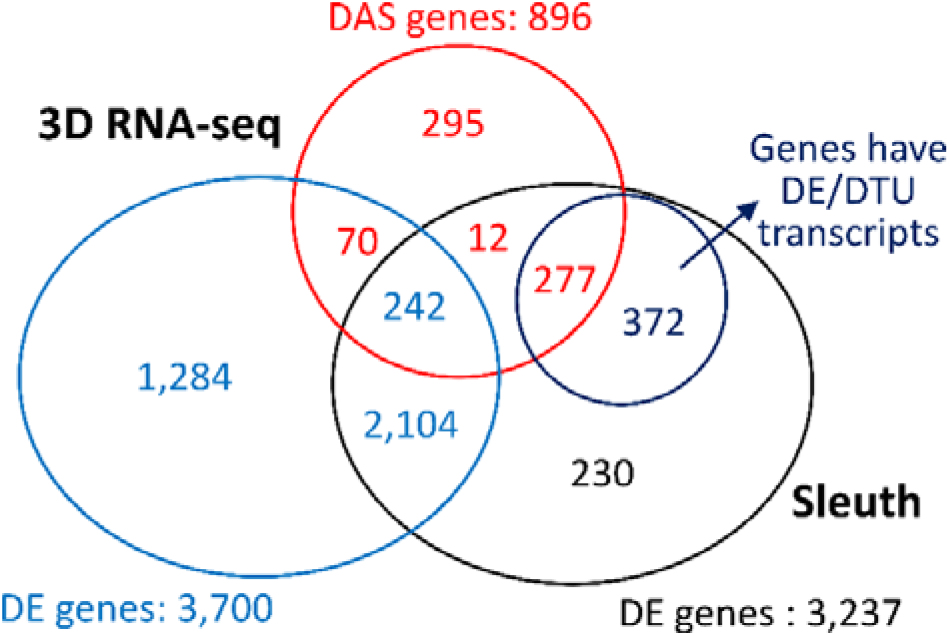
Comparison of the gene lists generated by 3D RNA-seq and Sleuth pipelines. The RNA-seq data on dexamethasone treatment of mice cells was taken from Frahm *et al*., (2018). Comparable parameters were applied when running 3D RNA-seq and Sleuth. The Venn diagram compares the DE genes from Sleuth to DE and DAS genes and DE and DTU transcripts from 3D RNA-seq.

**Figure 6.**
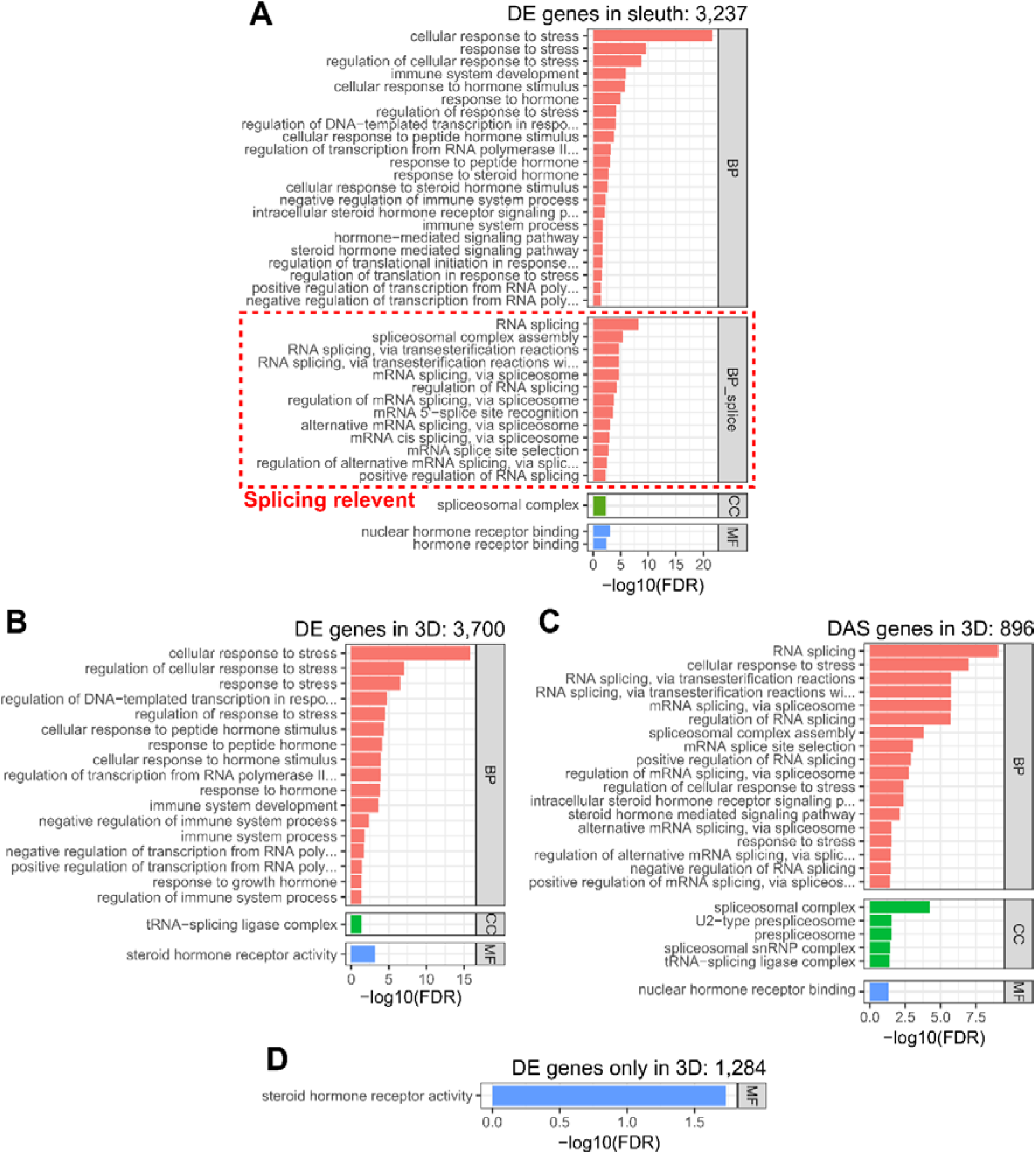
Top enriched GO terms identified from 3D RNA-seq and Sleuth. The Fisher’s exact test and topGO R package were used to generate significant enrichment gene ontology (GO) terms with FDR < 0.05. Significant terms relevant to response to stress, immune system, inflammation, hormone response, splicing/spliceosome are shown in the figure. A) Significantly enriched GO terms of DE genes from Sleuth; B) Significantly enriched GO terms of DE genes from 3D RNA-seq; C) Significantly enriched GO terms of DAS genes from 3D RNA-seq; and D) Significantly enriched GO terms of novel DE genes unique to 3D RNA-seq. Splicing/spliceosome related GO terms are enriched in the DE genes in *Sleuth* (red dashed box in A) but are found in GO terms associated with DAS genes in 3D RNA-seq (C). BP: Biological process; BP_splice: Biological process with terms of splice, splicing, spliceosome and spliceosomal; CC: Cellular Component; MF: Molecular Function.

The resolution of DE and DAS genes by *3D RNA-seq* was also illustrated in the GO and KEGG pathway enrichment annotations. The DE genes from *Sleuth* had significant terms and pathways that related to response to stress, response to hormone and immune system as well as multiple splicing/spliceosome terms indicating alternative splicing regulation of gene expression (Figure 6A; Additional file 1: Table S2A). The separation of DE and DAS genes by *3D RNA-seq* resolved the majority of stress, hormone and immune response terms to DE genes (transcriptional regulation) (Figure 6B; Additional file 1: Table S2B) and the splicing enrichment terms (such as spliceosome and mRNA surveillance pathways) to the DAS genes (AS regulation) (Figure 6C; Additional file 1: Table S2C). In addition, the 1,284 DE genes unique to the *3D RNA-seq* analysis (Figure 5) were enriched for steroid hormone receptor activity (Figure 6D) reflecting the nature of the chemical treatment.

The second comparison exploited the flexibility of *3D RNA-seq* to analyse the effects of multiple factors and detect sex-specific and brain region-specific DE and DAS genes and transcripts in the mouse data. In the original study, the effects of dexamethasone treatment on gene expression in NPSCs from different brain (cortical and hypothalamic) and sexes (male and female), were examined (Frahm et al., 2018). We therefore set up contrast groups (i.e. Female.Cortex.Dex vs Female.Cortex. Vehicle (untreated control), Male.Cortex.Dex vs Male.Cortex.Vehicle, Female.Hypothalmus.Dex vs Female.Hypothalmus.Vehicle and Male.Hypothalmus.Dex vs Male.Hypothalmus.Vehicle). The cut-offs were set as adjusted *p-value* <0.05, *L*_2_*FC* ≥ 1 and ΔPS ≥ 0.1. Across the four contrast groups, the 3D pipeline identified 930 DE genes and 509 DAS genes, of which 462 were only regulated by AS, and 2,121 DE transcripts and 628 DTU transcripts, of which 455 were AS regulated only. The results of individual contrast groups revealed that 1) both transcription and AS regulation were much more affected in cortex cells than in hypothalamic cells (Figure 7A-C) and 2) more genes and transcripts were down-regulated in cortex cells while in hypothalamic cells up-regulation dominated the expression changes (Figure 7A-C). The relative differences between brain regions at the DE gene level was described by Frahm *et al*., (Frahm et al., 2018). We also compared the DE genes from the 3D pipeline to those of Frahm *et al*., (Frahm et al., 2018) which used a *p-value* cut-off of < 0.05 (note: *p*-values were not adjusted and no *L*_2_*FC* cut-off was applied). There were 449 DE genes in common and 908 significant DE genes in Frahm *et al*., (Frahm et al., 2018) were filtered out in the 3D pipeline due to low *L*_2_*FC* or insignificant adjusted *p*-values (Figure 7D). Although stringent filters were used, the 3D analysis identified 481 unique DE genes. Finally, the isoform switch analysis identified 63 significant ISs of DAS genes in the contrast groups (Figure 7E). Thus, compared to the Sleuth pipeline, *3D RNA-seq* provides both transcript- and gene-level analysis to resolve both genes and transcripts differentially expressed due to transcriptional regulation, gene and transcript expression changes due to AS regulation by identifying novel DAS genes, DTU transcripts and ISs. The re-analysis of the mouse data with *3D RNA-seq* demonstrates how novel information can be unlocked from publicly available RNA-seq data which has only been used to analyse differential gene-level expression.

**Figure 7.**
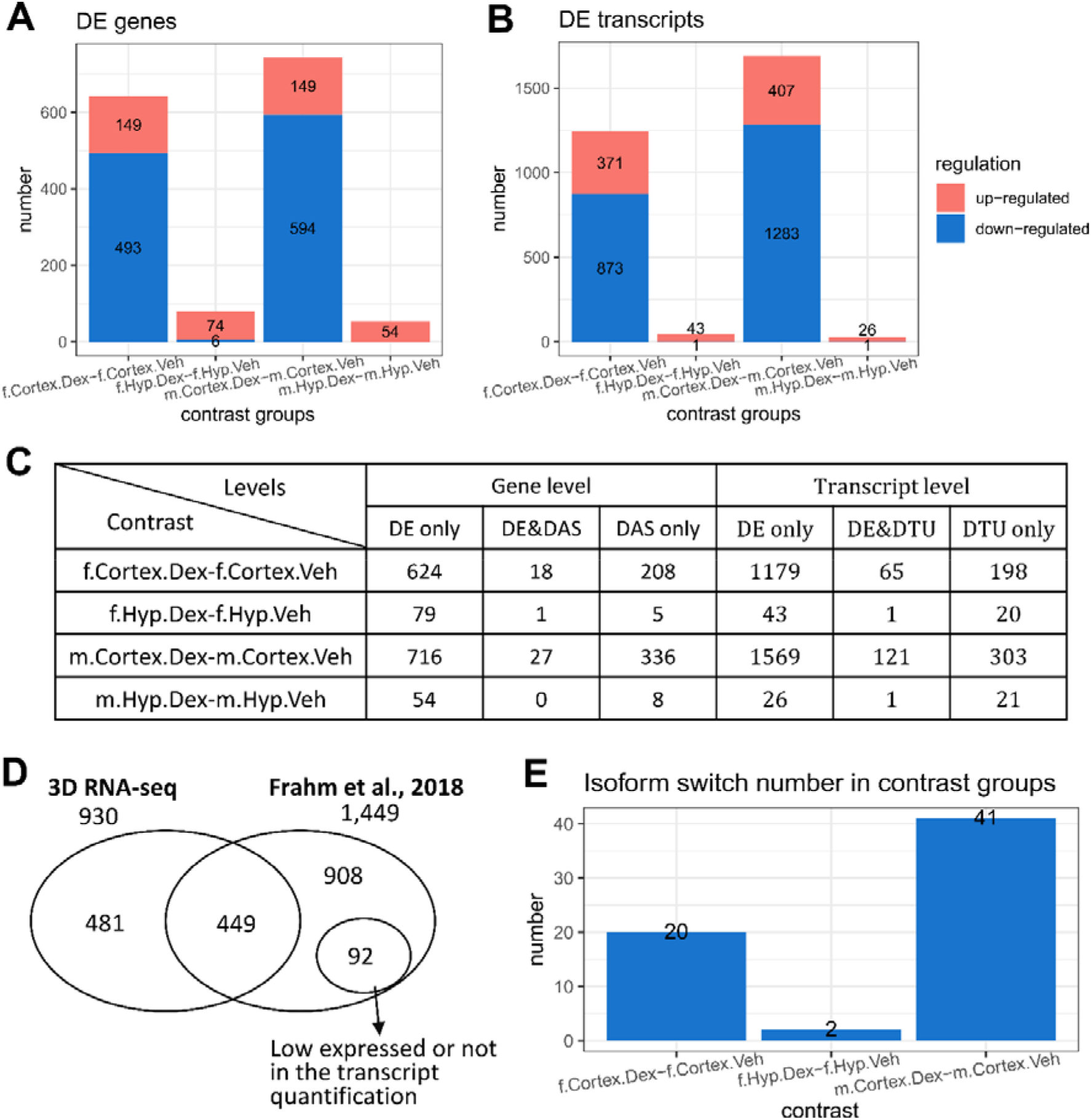
Sex-specific and tissue-specific expression analysis using 3D RNA-seq on the mouse data. Contrast groups were designed to investigate Dex-induced expression and alternative splicing changes between male and female and cortex and hypothalamus brain regions. Significant DE gene/transcript lists were generated by BH adjusted p-value < 0.05, *L*_2_*FC* ≥ 1 and Δ*PS* ≥ 0.1. A) Up- and down-regulated DE genes and B) DE transcripts. C) Summary of statistical analysis results from 3D RNA-seq in each contrast group. D) Venn diagram comparing the DE genes in the 3D RNA-seq analysis to the results in Frahm *et al*., (Frahm et al., 2018) in which the significant DE genes were determined by p-value < 0.05 (multiple testing adjustment and *L*_2_*FC* cut-off were not applied). 92 genes had low expression and were not included in the transcriptome quantification in 3D RNA-seq analysis. E) the number of Isoform switches in different contrast groups with the following cut-offs: probability of switch ≥ 0.5, difference of average TPMs at different conditions ≥ 1 TPM and adjusted p-value of the TPM difference < 0.05.

Finally, dexamethasone has recently gained prominence as it has been shown to reduce mortality in hospitalised patients with COVID-19 that require mechanical ventilation, supplementary oxygen, or Extra Corporeal Membrane Oxygenation (ECMO) (https://www.sps.nhs.uk/articles/dexamethasone-and-covid-19/). Our results raise the questions of whether dexamethasone treatment might affect processing of the viral RNA or alternative splicing of endogenous genes in patients. 3D RNA-seq would be a valuable tool investigating such questions.

## Discussion

### 3D RNA-seq is easy to use and designed for the maximum take-up by biologists

3D RNA-seq has the potential to revolutionise transcriptomics analysis by opening up RNA-seq analysis to wet lab scientists/biologists with little or no bioinformatics experience. The *3D RNA-seq* web-based application requires no installation and no programming skills from users. The whole analysis can be accomplished entirely within a web browser. The pipeline consists of 24 single steps within seven tabbed panels. It integrates widgets that provide users with technical summaries of the research behind each step so that they can set the parameters appropriate for their studies. Recommended parameters are also given in the manual and are used as defaults. Tutorial and demonstration videos are available on YouTube (see Availability of data and materials). More importantly, *3D RNA-seq* integrates interactive visualization at every step of the analysis so that users can visualize the changes caused by modification of the parameters as they move through the pipeline. Interactive visualization not only helps the user to understand the method behind each step better, but it also enables users to explore the optimum settings for the analysis, and ultimately offers reassurance and robustness in the results.

*3D RNA-seq* also allows the analysis of RNA-seq data with simple and complex experimental designs, such as time-series and developmental series data etc. The comparisons (contrast groups) can be set up between any pairs or groups of samples to cater for different investigations, a feature that is important for the universal applicability of *3D RNA-seq* for RNA-seq analysis. The *3D RNA-seq* App can be used to analyse RNA-seq data from any species for which transcript quantifications using *Salmon/Kallisto* can be generated. The accuracy of the quantifications using Salmon and Kallisto will depend on the quality and comprehensiveness of the reference transcriptome used to generate the transcript quantifications (Brown et al., 2017; Zhang et al., 2015, 2017). Multiple factors such as missing transcripts, transcript redundancy, transcript fragments and mis-annotations at the 5’ and 3’ ends of isoforms can drastically affect accuracy of transcript quantification (Alamancos et al., 2015; Soneson et al., 2019; Zhang et al., 2015, 2017). For many species, comprehensive, optimised transcriptomes do not exist and using partial, incomplete transcriptomes and/or which have not been filtered to remove such erroneous isoform information will also produce inaccurate transcript and AS quantification. As transcriptomes improve for many species, transcript quantification and results from *3D RNA-seq* will also improve.

The speed and accuracy of the *3D RNA-seq* analysis has the potential to revolutionise gene expression research programmes. Generation of transcript quantifications and running *3D RNA-seq* is performed in less than three days. It reduces an analytical step which previously required skilled bioinformaticians and often took many weeks or even months to complete. This is significant not only in terms of satisfaction in result generation for the individual scientists/biologists but also for strategic planning of research programmes where now multiple, consecutive RNA-seq experiments can be planned within the period of funding proposal. More importantly, the rapid turn-over of the analysis provided by *3D RNA-seq* creates a level playing field for research groups with very different access to bioinformatics expertise or resources thereby supporting, in particular, smaller groups and early career scientists. *3D RNA-seq* will also aid research projects in developing countries with limited access to computational infrastructures and bioinformatics expertise. Although the quantification of transcripts is not included in 3D RNA-seq, we have provided a tutorial document and Youtube video on performing such tasks using publicly available Galaxy servers (https://usegalaxy.eu/) (see Methods and Accessibility of data and materials). The transcript quantification output from Galaxy can be used as input to 3D RNA-seq directly.

### 3D RNA-seq facilitates speedy publication of RNA-seq analysis and improves the transparency and reproducibility of the analysis

The fast turn-over of the analysis provided by *3D RNA-Seq* will present a significant advantage for speedy publication of the results. In *3D RNA-seq*, multiple types of figures/summaries are generated and saved including commonly used visualizations such as heat-maps, GO enrichment plots, expression profile plots, volcano plots etc. Users can easily generate and save new figures for each new set of parameters. The format, resolution and size of the figure can be customised and previewed in *3D RNA-seq* before saving. In addition, the significant DE/DAS/DTU/IS gene and transcript lists are saved for further interpretation. The final technical report generated in the final step of *3D RNA-seq* records each step in the analysis with the parameters used and integrates all the saved figures. The report is comprehensive, accurate and reproducible including all the information required for “Material and Methods” sections for the RNA-seq analysis as well as figures for the Results and Supplementary Materials. Furthermore, some publications have very poor technical descriptions of the methods and parameters used for RNA-seq analyses. In future, submission of such a report along with manuscripts to scientific journals would facilitate transparency and allow reviewers to identify issues with the analysis and how it relates to other published work.

### 3D RNA-seq provides enhanced alternative splicing analysis at the transcript level

Over the last 20 years, genome-wide expression analyses have relied on microarrays and more recently, RNA-seq analyses mostly at the gene level. RNA-seq allows gene expression to be analyzed at the level of both genes and transcripts which in turn provides a means to study post-transcriptional processes such as alternative splicing. AS regulation plays key roles in gene expression and novel genes with altered gene expression at the AS level have been found in most of the RNA-seq studies. Despite the importance of AS and the potential to include AS analysis routinely in RNA-seq, AS is still largely ignored. For example, 4,065 publications from the Web of Science Core Collection (2008-2019) (https://wok.mimas.ac.uk/) were retrieved with the term “RNA-seq differential gene expression”, but only ca. 289 included “differential alternative splicing”. *3D RNA-seq* provides by far the most detailed differential expression analysis on the transcript and alternative splicing level. In particular, it allows the identification of both DAS genes and DTU transcripts which are differentiated from DE genes/transcripts and a range of measurements, visualizations and statistical tests provide a comprehensive analysis of alternative splicing alongside differential expression. Additionally, *3D RNA-seq* has also integrated methods that detect isoform switches which can play pivotal roles in re-programming of gene expression through switching of functionally different transcript isoforms between, for example, normal and tumor tissues to provide signatures for cancer diagnostics and prognostics (Sebestyén et al., 2015). With the enhanced AS analysis in *3D RNA-seq*, key and novel genes under AS regulation which underpin important biological processes can be identified and provide new targets for medical intervention or crop improvement.

### 3D RNA-seq unlocks the discovery potential for RNA-seq data

Finally, the speed of *3D RNA-seq* now makes it feasible for biologists to re-analyse existing or publicly available RNA-seq data to give improved differential expression analysis and novel AS information as demonstrated here for the mouse data. Datasets from different labs can now be quickly compared using the same parameters and without performing new experiments. In addition, with new transcriptome releases, it is feasible to re-analyse datasets and update results. *3D RNA-seq* provides a reproducible platform to standardise analyses and utilise new and existing data for improved resolution and interpretation.

## Materials and Methods

We have developed 3D RNA-seq App, an R package which provides a web-based shiny App for flexible and powerful differential expression and alternative splicing analysis of RNA-seq data. The program is designed for ease-of-use and can be run by biologists with minimal bioinformatics experience or by bioinformaticians with little exposure to RNA-seq analysis.

The whole analysis from transcript quantification to data pre-processing, differential gene and transcript expression, differential alternative splicing, differential transcript usage and isoform switching analysis can be performed in less than three days. The analysis is optimized with quality control plots and generates a report containing customisable publication-quality figures and data tables to aid interpretation of results.

3D RNA-seq enables routine analysis of differential alternative splicing and isoform switching and can handle complex experimental designs. The speed and accuracy of the analysis has the potential to revolutionise what is achievable with RNA-seq technology.

### Input files for 3D RNA-seq App

Three simple input files are required for *3D RNA-seq* analysis (Additional file 1: Figure S5). 1) A meta-data table in “csv” (comma delimited) format containing the information of experimental design, including conditions/treatments, biological replicates, sequencing replicates (same biological sample divided and sequenced on different sequencing lanes) and quantification file names (Additional file 1: Figure S5A). 2) Transcript quantification outputs generated using *Salmon* (Patro et al., 2017) or Kallisto (Bray et al., 2016) (Additional file 1: Figure S5B). Transcript quantification is the first difficulty for most of the biologists who are lack of programming skills. A detailed user manual with demo video of quantification by using Salmon/Kallisto on the Galaxy interface (*Galaxy | Europe*, n.d.) has been provided (see Availability of data and materials). 3) A transcript-gene association file consisting of a “csv” spreadsheet with the first column listing transcript names and second column listing gene IDs. This relates transcript names to gene IDs which allows expression analysis at both gene and transcript level and the AS analysis within each gene (Additional file 1: Figure S5C). It is also possible to extract this info from other files when that information is available (Additional file 1: Figure S5D and E). All the inputs can be easily uploaded by selecting corresponding files from a local computer.

### Data pre-processing with optimal quality control

First, *Salmon* and *Kallisto* outputs (“quant.sf” from *Salmon* and “abundance.tsv” from *Kallisto*) for each sample, will be imported into *3D RNA-seq* and read counts will be generated using the tximport R package (Soneson et al., 2016), which takes into account the transcript length and library size. Sequencing replicates, if present, will be merged to increase sequencing depth. Then, the data is pre-processed in various steps to reduce noise (e.g. removal of low expressed genes and transcripts) and technical variance (e.g. batch effects). In each step, interactive visualization plots are produced to show the immediate impact of the parameter change and facilitate the optimization of parameters for pre-processing. 1) Removal of low expressed transcripts. RNA-seq read counts follow approximately the negative binomial distribution. The variance of log_2_ transformed read counts decreases monotonically with the increase of mean (Additional file 1: Figure S2) (Benaroya & Mi Han, 2005; Law et al., 2014; McCarthy et al., 2012). However, low expressed transcripts do not follow such a distribution and thus should be removed. On the basis that an expressed transcript should have a minimum count per million (CPM), *n*, in at least *m* samples, the cut-off can be adjusted by the user and the effect on mean-variance trends visualised immediately. 2) Similarities and differences between samples are visualized by Principal Component Analysis (PCA) plots (Additional file 1: Figure S3). PCA is a method to project data variance of thousands of variables (transcripts/genes) to a few principal components (PC) that capture the majority of the data variability. PCA plots can reveal major variances in an experiment in an unbiased manner which allows technical effects (e.g. batch effects among biological replicates caused by different experimental conditions) to be identified. 4) Correction of technical effects if required. Batch effects, if detected, can be corrected by different methods such as the RUVSeq R package, which estimates a co-efficient for batch effect term which can be incorporated into the design matrix with the main factors in linear regression models for 3D analysis (Additional file 1: Figure S3B) (Risso et al., 2014). 5) Normalization of read counts across the libraries. Read counts can be normalized using Trimmed Mean of M-values (TMM), Relative Log Expression (RLE) or upper-quartile method (Bullard et al., 2010). Read count distributions before and after normalization can be visualized in the plots.

### Principle of 3D analysis: identification of DE, DAS and DTU genes and transcripts

RNA-seq experimental designs often involve a single, or multiple experimental conditions that may affect gene expression. The *3D RNA-seq* App provides a flexible way to set up contrast groups where users can select any samples or groups of samples of interest for comparison. Thus, it allows analyses of both simple pair-wise comparisons and complex experimental designs such as time-series, developmental series and multiple conditions (Additional file 1: Figure S4). For each contrast group, different statistics are defined for robust DE/DAS/DTU (3D) predictions (Figure 1). 1) For differential expression, log_2_ fold change (*L*_2_*FC*) is the difference of log_2_-CPM values in contrast groups; 2) for differential alternative splicing, Δ percent spliced (Δ*PS*) is the difference of *PS* values which are defined as the ratios of transcript average abundances divided by the average gene abundances; and 3) *p*-values of multiple comparisons are adjusted to control the false discovery rate (FDR) (Yoav Benjamini & Yekutieli, 2001).

Stringency of the analysis can be modified by changing the cut-off settings. A gene/transcript is identified as DE in a contrast group if *L*_2_*FC* of expression is greater than or equal to an established cut-off value (e.g. *L*_2_*FC* ≥ 1) and with adjusted *p*-value less than a cut-off (e.g. *p*<0.01 or 0.05) (Figure 1). At the AS level, gene expression is compared to transcript expression in the contrast groups (Ritchie et al., 2015). To identify DAS genes, the expression of each transcript is compared to the weighted average expression of all the transcripts for the same gene (weight on transcript expression variance). The *p*-value of each test is converted to gene-wise *p*-value by using the F-test or Simes method (Ritchie et al., 2015). To identify DTU transcripts, each transcript is compared to the weighted average of all the other transcripts of the same gene (Figure 1). A gene is DAS in a contrast group if the adjusted *p*-value is less than an established cut-off value and at least one transcript of the gene has a Δ*PS* greater than an established cut-off value (e.g. Δ*PS* >0.1). A transcript is DTU if the adjusted *p*-value and absolute values of ΔPS is greater than the selected cut-off values, respectively. *3D RNA-seq* thereby identifies significant DE genes and transcripts, DAS genes and DTU transcripts and these results are saved in summary figures (see Figure 3A and B), and tables of identified transcript/gene lists in csv files.

### 3D RNA-seq is flexible and can work with all experimental designs

Within 3D RNA-seq, the contrast groups (comparisons) can be set-up in multiple ways: 1) between any two samples with biological replicates not just limited to the comparison of two experimental control samples only; 2) between groups of samples, where the samples can be grouped at a higher level (e.g. group samples by gender); 3) contrast groups can be set up to test the interactions of different factors (e.g. the interactions between gender and treatment. Do different genders respond to the treatment differently?); 4) Besides discrete comparisons of samples and groups, 3D RNA-seq also allows the comparison of trend changes over time in each parallel condition group by using the spline method (Hastie & Chambers, 1992) to study dynamic activities and changes of transcriptome over time and over several conditions simultaneously. For example, the identification of the dynamics of stress-responsive genes and AS events in plants (Bahieldin et al., 2015; Calixto et al., 2018, 2019; Ma et al., 2019), timing mechanisms of circadian clocks (Blair et al., 2019; J. A. Kim et al., 2019; Oh et al., 2014) and drug responses (Jardim-Perassi et al., 2019; Kang et al., 2019). Adjusted p-values are used to report the genes and transcripts that show significant differences between samples/groups/series at both levels of transcription and AS (DE, DAS and DTU).

### Transcript isoform switch analysis

Transcript isoform switches (ISs) are a prominent type of DAS within a gene where a pair of alternatively spliced isoforms reverse the order of their relative expression abundances in response to stimulus (Guo et al., 2017). In the *3D RNA-seq* analysis pipeline, the iso-kTSP method is introduced to study the ISs between pair-wise conditions in the user-defined contrast groups (Sebestyén et al., 2015) while the Time-Series Isoform Switch (TSIS) method is used to identify the ISs between any consecutive time-points in time-series or developmental data (Guo et al., 2017).

### Outputs and visualization

*3D RNA-seq* allows users to save results of significant DE/DAS/DTU gene/transcript lists, intermediate data of the whole analysis and publication-quality plots to a local folder by clicking “action” buttons at the various steps in the App. Four folders are created to save 3D analysis outputs, “result”, “figure”, “report” and “data”. The gene and transcript expression in read counts and TPMs, testing statistics and analysis results of 3D analysis will be saved as “.csv” files in the “result” folder. All the figures of data pre-processing, 3D analysis and downstream visualization are saved as “png” and “pdf” formats with user specified width, height and resolution in the “figure” folder. The intermediate variables generated during 3D analysis are saved as “.RData” R objects in the “data” folder for R users to carry out further analysis if required. In the last step of 3D analysis, a report in three formats, “html”, “pdf” and “word”, will be generated in the “report” folder using *R Markdown* (Baumer & Udwin, 2015). The report includes sections of “Methods”, “Results”, “Supplementary figures”, “Supplementary material” and “References”, in which all the parameters selected by the users for each step are recorded. The publication-quality figures and reports provide all information required for publications and it also provide sufficient information to reproduce the same analysis and results.

Genes and transcripts that are significantly changed between contrast groups can be visualized within the App in Venn diagrams, histograms and volcano plots (e.g. the number of up- and down-regulated DE genes and transcripts, isoform switches etc.) (see Figure 3C-E). Heat-maps are used to visualize co-expression clusters of significant 3D genes and transcripts and investigate their expression pattern changes across conditions (Figure 4A and B). Individual gene and transcript profiles and ΔPS plots can be generated to give users intuitive visualization of those with significant abundance and AS changes, thereby allowing a convenient and detailed investigation of individual genes/transcripts and providing a means of selecting candidate genes and transcripts for experimental validation and future research. The lists of genes and transcripts with significant changes in DE/DAS/DTU can be downloaded and exported out of 3D RNA-seq for gene ontology (GO) enrichment analysis using appropriate web databases or GO analysis programs. The significantly enriched annotation results can be re-imported into the *3D RNA-seq* App to generate GO annotation plots (Additional file 1: Figure S6 and see Figure 4C and D). To generate publication-quality figures, users can set widths, heights, resolution and colours for histograms, plots, heat-maps etc. to be saved. The analysis reports contain full details of the custom configured parameters, methods and results tables and figures from the entire 3D analysis. The analysis report is a detailed document that guarantees the reproducibility of the analysis and results.

### 3D RNA-seq adopts the best practice and integrates the state-of-art methods for DE and DAS analysis

*3D RNA-seq* adopts best practice and integrates state-of-art methods for RNA-seq data pre-processing: 1) Tximport to convert TPM values to read counts while taking transcript length and sequencing depth into account (Soneson et al., 2016); 2) mean-variance trend plots to filter low expressed genes/transcripts to improve fit to statistical models and remove discrepancies due to read count distribution assumptions (Law et al., 2014); 3) PCA plots to visualize sample variances and identify technical effects; 4) RUVSeq to estimate batch effects (Risso et al., 2014); 5) a number of read count normalization methods to correct the variances and reduce the false positives for highly abundant transcript outliers (Bullard et al., 2010).

*Limma* voom (Law et al., 2014) was chosen as the engine for the DE analysis (DE, DAS and DTU) for five reasons. Firstly, from different studies, *Limma* is consistently one of the best performing methods for RNA-seq differential expression analysis and has a good control of FDR (Pimentel et al., 2017; Rapaport et al., 2013; Tang et al., 2015). Secondly, it allows both the DE and DAS analysis within the same framework. More importantly, the DE analysis and DAS analysis differentiates genes that are affected by transcriptional regulation (e.g. by transcriptional factors) and alternative splicing regulation (e.g. by splicing factors) as shown in Figures 3-7. Thirdly, *Limma* employs a linear model that runs very quickly and compared with methods requiring bootstrapping data, such as Sleuth (Pimentel et al., 2017), the running time and memory required is significantly reduced. Fourthly, *Limma* has good compatibility with other bioinformatics R packages for a range of other functions, such as estimation of batch effects and normalisation of sequencing biases. In addition to batch effects that occur systematically for all the samples in a certain biological replicate, RNA-seq data may have variations in sample quality due to, for example, degradation or contamination of specific samples. These problematic samples are often shown as outliers in the PCA plot. The *Limma*-voomWithQualityWeights function has been incorporated in the 3D RNA-seq App to balance the outliners (Law et al., 2014; Liu et al., 2015). Lastly, but most importantly, *limma* allows flexible experimental designs, where any pairs or groups of samples can be compared in contrast to most of the current DE tools which offer limited choices for comparisons.

### Availability of the 3D RNA-seq App

The *3D RNA-seq* App is available as a web service and an R package. 1) A publicly accessible version of *3D RNA-seq* is running at the James Hutton Institute and can be viewed at: https://3drnaseq.hutton.ac.uk/. The user can upload input files and carry out the entire analysis within a browser. The intermediate data and results of 3D analysis can be downloaded during the final step. This web interface allows biologists to use *3D RNA-seq* without installing it. The *3D RNA-seq* manual and a YouTube tutorial (https://www.youtube.com/watch?v=rqeXECX1-T4) to demonstrate the usage of 3D RNA-seq and upstream quantification step using Salmon can be found at the same web address; 2) The R package *ThreeDRNAseq* can be installed locally and users can run the *3D RNA-seq* App by typing a command “run3DApp()” through RStudio on a local PC. This version is suitable for people with a good knowledge of R and who prefer to run the analysis locally. The user manuals for both using 3D RNA-seq App and command lines of ThreeDRNAseq R package for 3D analysis is provided at: https://github.com/wyguo/ThreeDRNAseq/tree/master/vignettes/user_manuals

## Acknowledgements

We thank Craig Simpson (The James Hutton Institute), Vivek Raxwal (Masaryk University, Czech Republic), Pingtao Ding (The Sainsbury Laboratory, Norwich) for testing the *3D RNA-seq* App.

## Additional information

### Availability of data and materials

The *3D RNA-seq* web interface is available at https://3drnaseq.hutton.ac.uk. The R package version (ThreeDRNAseq) is available on Github at https://github.com/wyguo/ThreeDRNAseq. Manuals for both versions and transcript quantification on Galaxy interface can be accessed from https://github.com/wyguo/ThreeDRNAseq/tree/master/vignettes/user_manuals Tutorial and demo video can be viewed from https://www.youtube.com/watch?v=rqeXECX1-T4

The *Kallisto* transcript quantifications from the dexamethasone treatment on mice were downloaded from:

https://figshare.com/articles/kallisto_quantifications_of_Frahm_et_al_2017/6203012.

The Sleuth/aggregated *p*-values pipeline is at: https://pachterlab.github.io/sleuth_walkthroughs/pval_agg/analysis.html.

### Authors’ contributions

RZ and JB supervised all aspects of the work; WG, RZ, JB designed the 3D analysis pipeline and WG coded the whole 3D RNA-seq App; NT and CC collected and prepared the Arabidopsis cold response RNA-seq data; GS and IM set up and maintained the web service; WG, RZ JB and RW drafted the manuscript; All authors engaged in discussions and testing of the 3D RNA-seq App; All authors were involved in revision and improvement of the final manuscripts; RZ, JB and RW interpreted the main findings.

## Additional file 1

### Supplementary Figures

**Figure S1:**
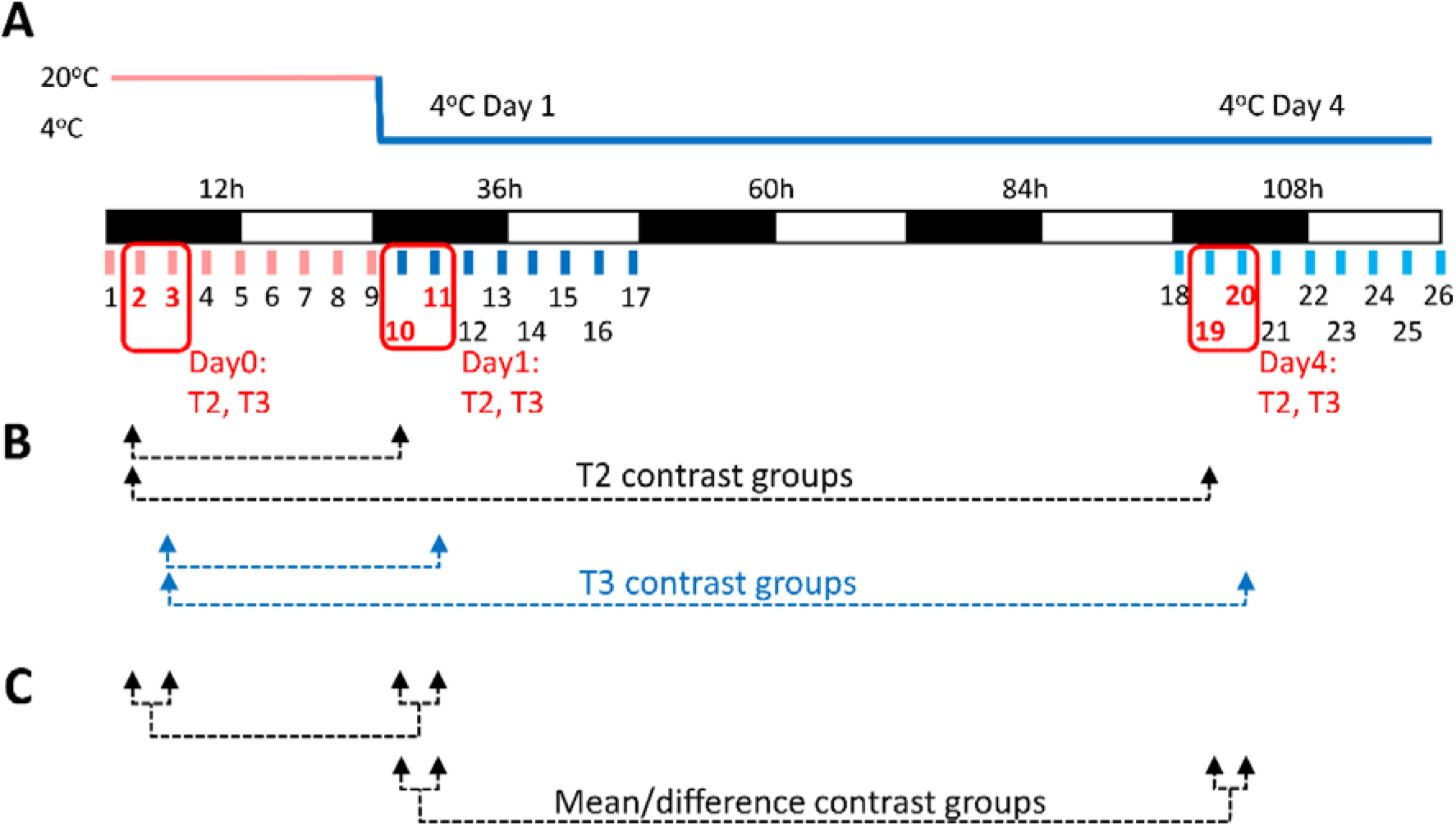
Example dataset and flexibility of the analysis. A) Data from six time-points (T2, T3 of 20°C Day, T10, T11 of Day 1 at 4°C, T19, T20 of Day 4 at 4°C) from a time-series of Arabidopsis plants exposed to cold (Calixto et al., 2018) were used as example data to illustrate setting up of contrast groups and analysis. B) Setting up the contrast groups between the same time points in different days to look for the effect of the duration of cold stress. C) Setting up the contrast groups between the mean of multiple groups or difference between pair-wise groups.

**Figure S2:**
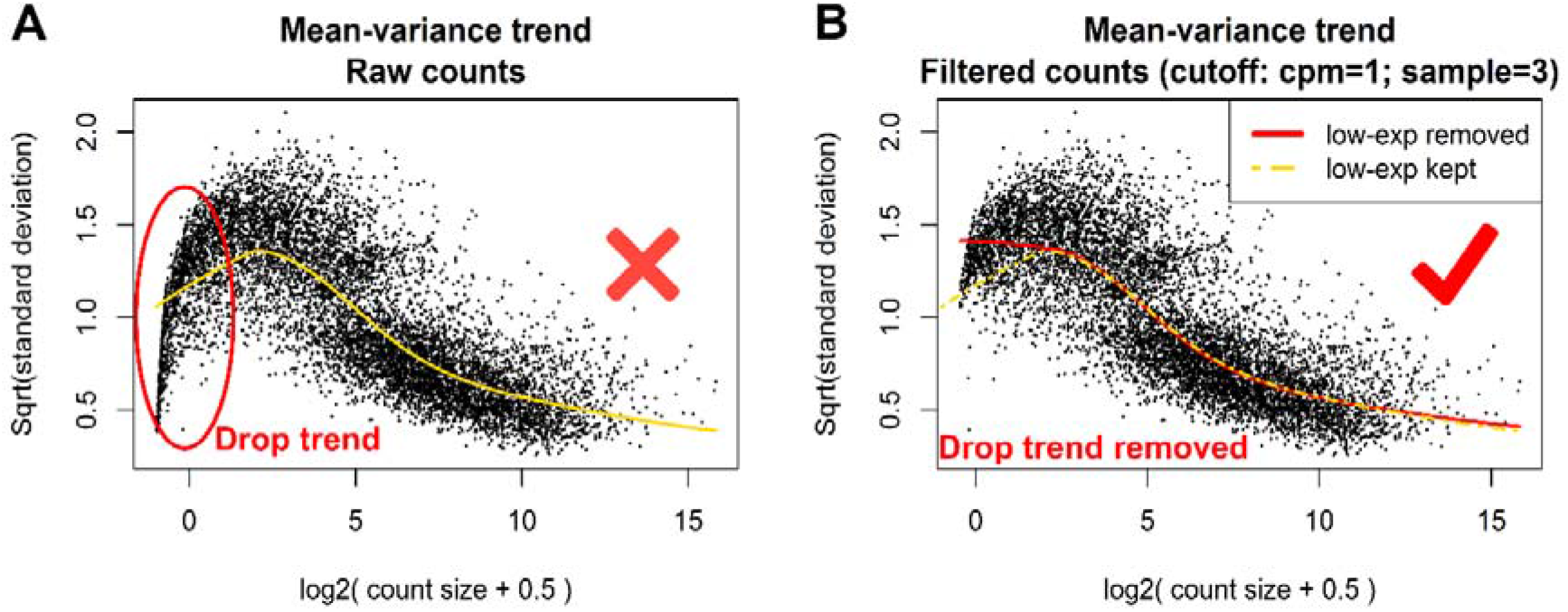
Filtering of low expressed transcripts based on mean-variance trend. A) and B) are the mean-variance trend plots before and after filtering low expressed transcripts using Arabidopsis cold response data, respectively. In the plots, each black point represents a transcript. The red and yellow curves are the fitted trends of these points. The red circle in plot A) highlights the drop trend of low expressed transcripts.

**Figure S3:**
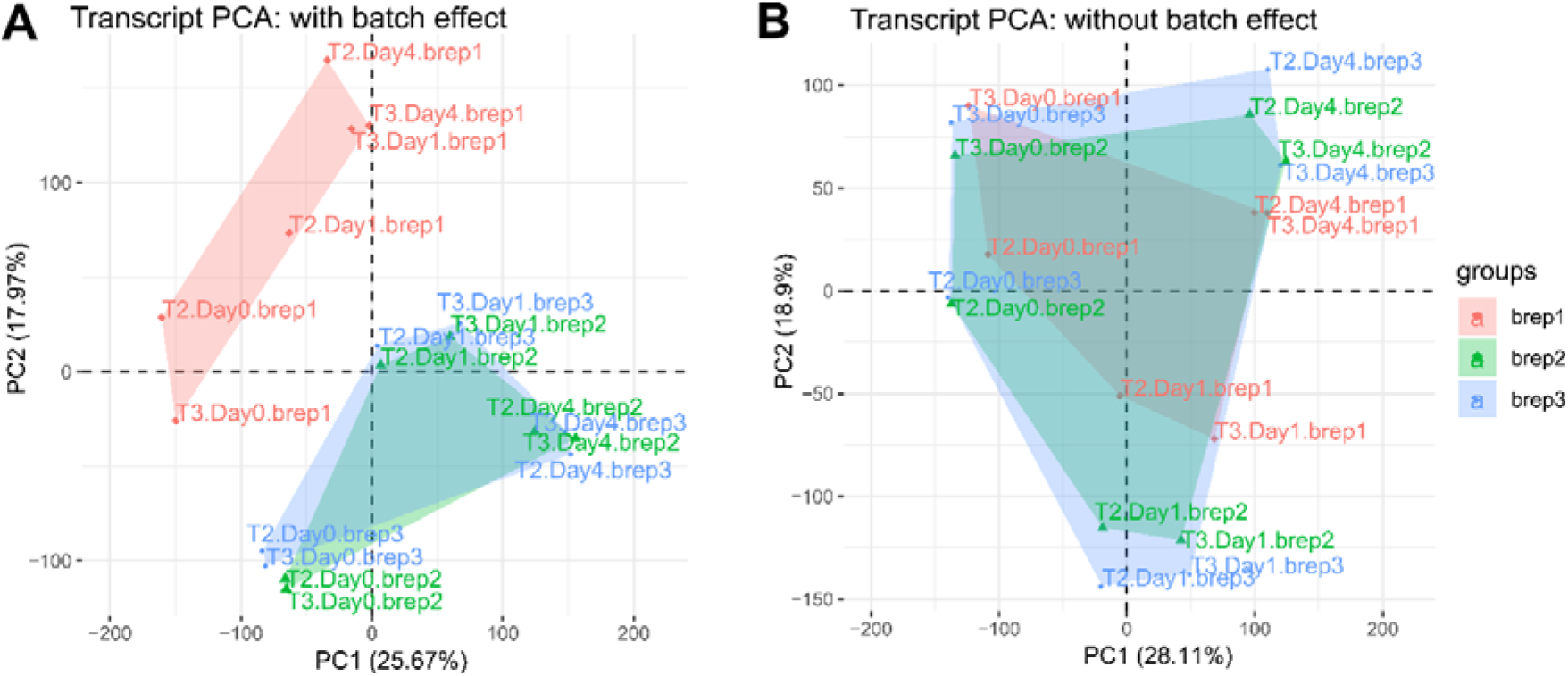
PCA plots of samples from Arabidopsis cold response data based on transcript level expression. A) No adjustment of batch effects. B) Batch effects of biological replicates were removed from the data using the RUVSeq method.

**Figure S4:**
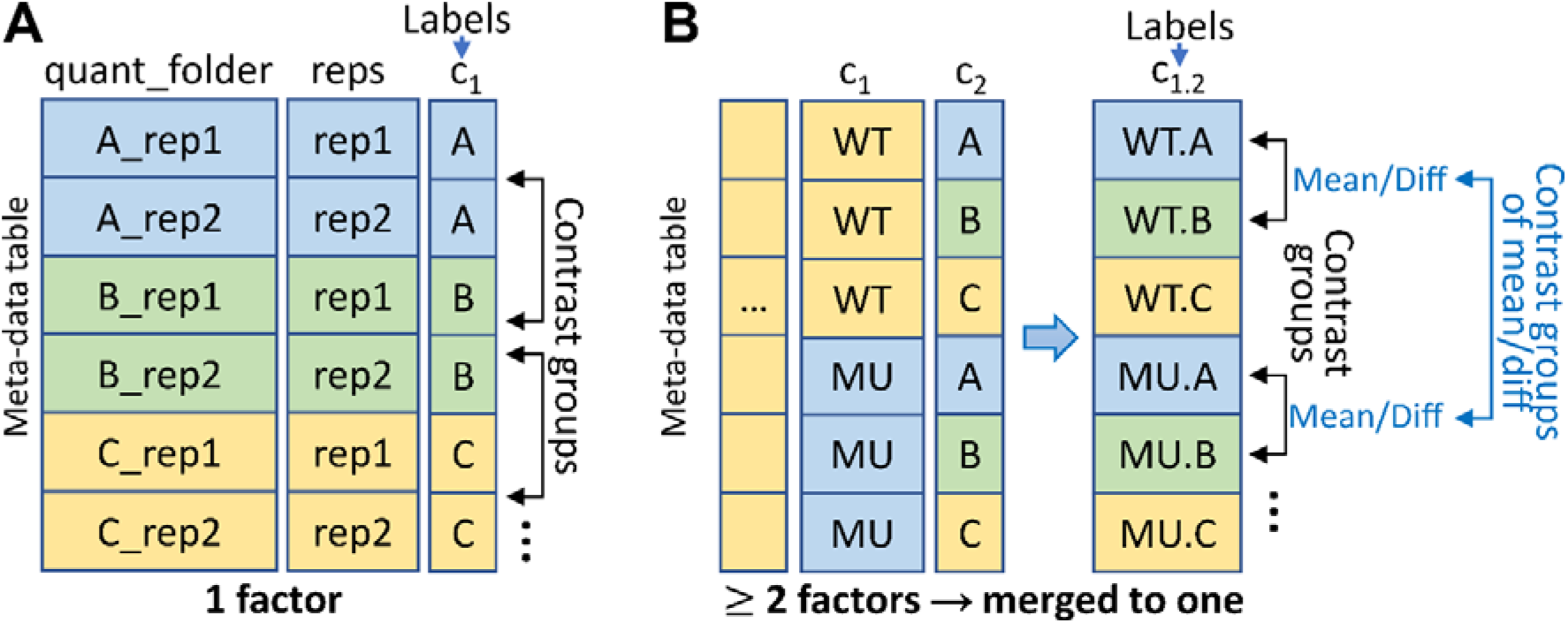
Generation of labels and setting contrast groups for comparisons. Meta-table of A) single factor and B) multiple factors. In the meta-data table of the experimental design (Supplemental Figure 1A), if multiple factors are selected, they will be combined to create a single column of grouping labels. 3D RNA-seq provides a flexible way where users can generate contrast groups of simple pair-wise analyses and complex experimental design such as time-series, development series and multiple conditions from grouping labels. For example, in B), which compares wild-type (WT) and mutant (MU) under three different conditions, contrast groups can be set as WT.B-WT.A (WT.B vs WT.A) and MU.B-MU.A (MU.B vs MU.A) to compare expression of condition WT.B to WT.A and MU.B to MU.A, respectively. In addition, users can set contrast group (MU.B+WT.B)/2-(MU.A+WT.A)/2 to compare the mean of multiple conditions and set contrast group (MU.B-WT.B)-(MU.A-WT.A) to compare the differences of pair-wise groups to study the interactions between the genotype and the conditions.

**Figure S5:**
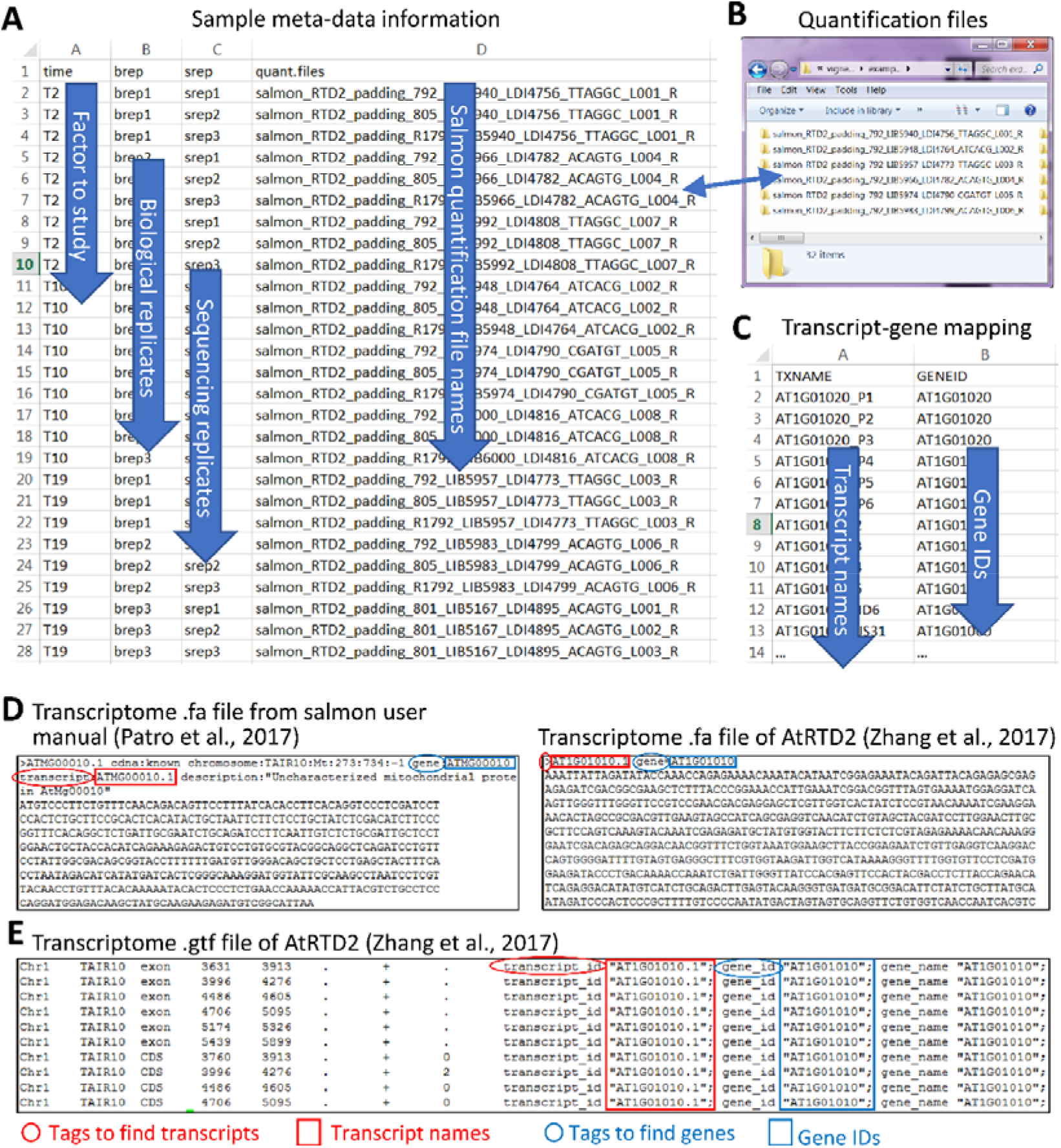
Input files of 3D RNA-seq analysis. A) A meta-data “csv” file containing all the experimental information for each sample. B) A folder of transcript quantification files from Salmon (Patro et al., 2017). C) A transcript-gene association mapping “csv” file. Alternatively, users can provide D) the transcriptome “fa” or E) “gtf” files to the App to generate the transcript-gene association information. The transcript names and gene IDs will be extracted from specific tags.

**Figure S6:**
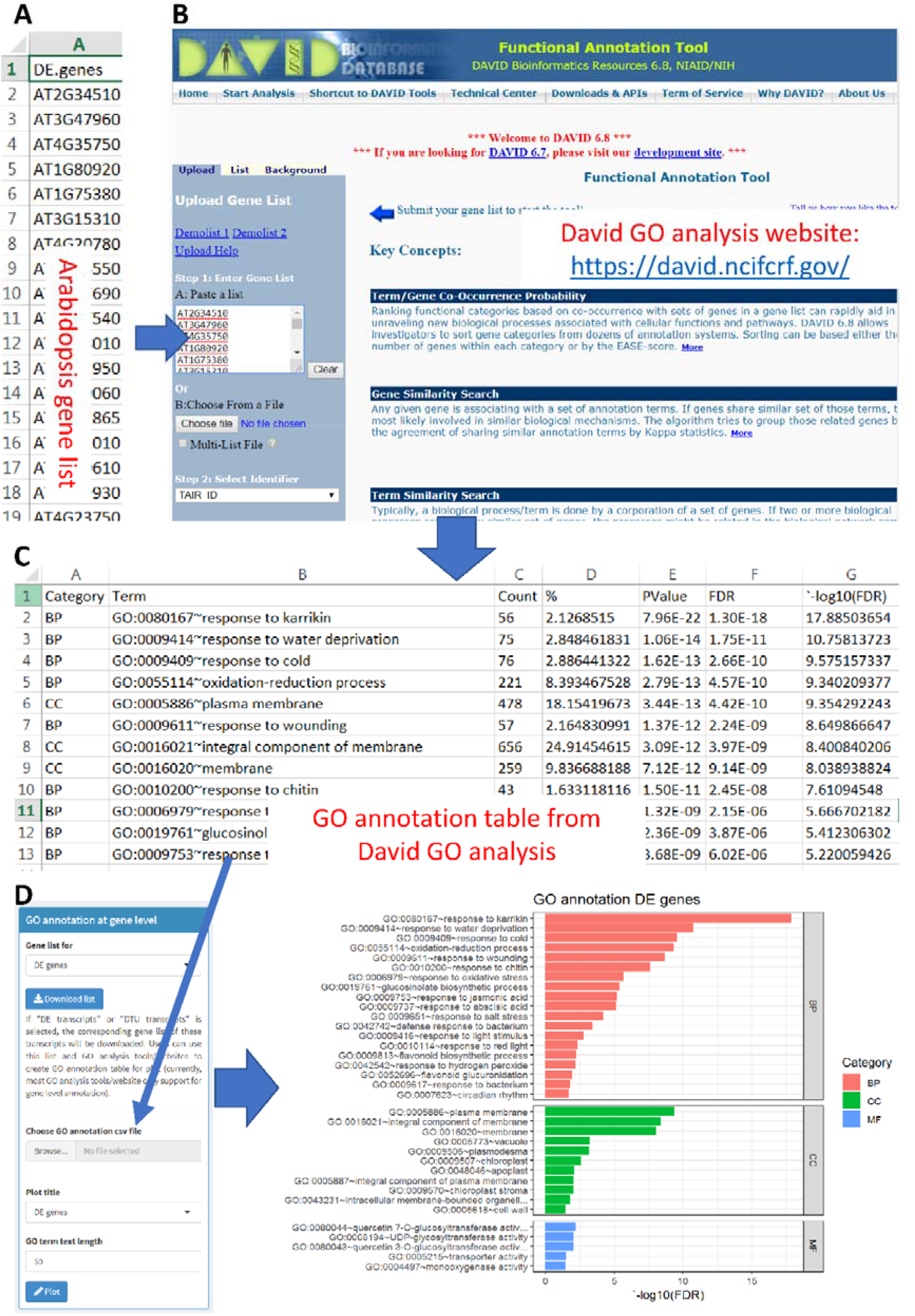
Process for making GO annotation plots. To generate GO annotation plots, A) the gene lists of DE genes, DAS genes, DE transcripts and DTU transcripts generated by 3D RNA-seq are downloaded as a spreadsheet; B) A gene list is selected and uploaded to functional annotation tools/website for GO analysis, e.g. the David https://david.ncifcrf.gov/ and agriGO http://bioinfo.cau.edu.cn/agriGO/ for plant species; C) the significantly enriched GO terms of genes are saved to a csv file (comma delimited) and D) the csv file of GO terms is uploaded to 3D RNA-seq App to make the GO annotation plot.

### Supplementary Tables

**Table S1:**
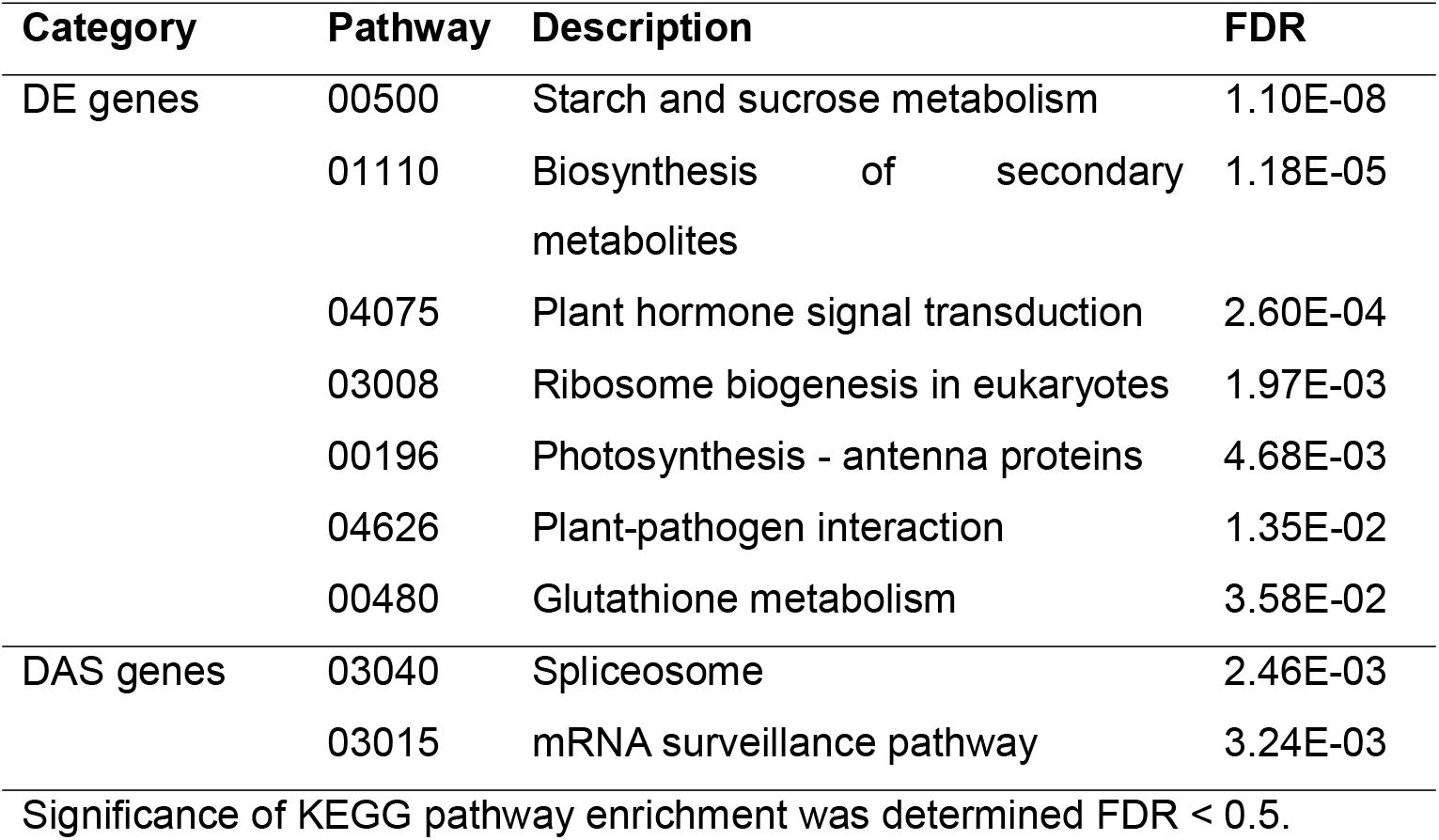
KEGG pathway enrichment analysis results of the Arabidopsis data.

**Table S2:** KEGG pathway enrichment analysis results of the mouse data.

| Category | Pathway | Description | FDR |
| --- | --- | --- | --- |
| **A**<br>3,237 DE genes<br>from Sleuth | 04919 | Thyroid hormone signalling pathway | 2.87E-06 |
|  | **03040** | **Spliceosome** | **5.46E-06** |
|  | 04935 | Growth hormone synthesis, secretion and action | 2.51E-05 |
|  | 05418 | Fluid shear stress and atherosclerosis | 3.33E-05 |
|  | 05206 | MicroRNAs in cancer | 8.73E-05 |
|  | **03015** | **mRNA surveillance pathway** | **5.72E-04** |
|  | 04928 | Parathyroid hormone synthesis, secretion and action | 9.33E-04 |
|  | 03013 | RNA transport | 5.53E-03 |
|  | 00970 | Aminoacyl-tRNA biosynthesis | 5.92E-03 |
|  | 04918 | Thyroid hormone synthesis | 1.95E-02 |
|  | 03018 | RNA degradation | 4.61E-02 |
| **B**<br>3,700 DE genes<br>from 3D RNA-seq | 05418 | Fluid shear stress and atherosclerosis | 7.28E-06 |
|  | 04919 | Thyroid hormone signalling pathway | 4.62E-04 |
|  | 04935 | Growth hormone synthesis, secretion and action | 2.72E-03 |
|  | 03018 | RNA degradation | 1.80E-02 |
|  | 04928 | Parathyroid hormone synthesis, secretion and action | 1.83E-02 |
|  | 03013 | RNA transport | 3.89E-02 |
| **C**<br>896 DAS genes<br>from 3D RNA-seq | **03040** | **Spliceosome** | **3.52E-05** |
|  | 04919 | Thyroid hormone signalling pathway | 3.94E-03 |
|  | 03013 | RNA transport | 8.10E-03 |
|  | 00970 | Aminoacyl-tRNA biosynthesis | 3.49E-02 |
|  | 05418 | Fluid shear stress and atherosclerosis | 4.15E-02 |
|  | **03015** | **mRNA surveillance pathway** | **4.15E-02** |

Only the significant pathways (FDR < 0.5) relevant to response to stress, immune system, inflammation, hormone response, splicing/spliceosome are shown in the table.

